# Opposing effects of population density and stress on *Escherichia coli* mutation rate

**DOI:** 10.1101/256305

**Authors:** Rok Krašovec, Huw Richards, Danna R. Gifford, Roman V. Belavkin, Alastair Channon, Elizabeth Aston, Andrew J. McBain, Christopher G. Knight

**Affiliations:** Faculty of Biology, Medicine and Health, The University of Manchester, Manchester, M13 9PT, United Kingdom; Faculty of Science and Engineering, The University of Manchester, Manchester, M13 9PT, United Kingdom.; School of Engineering and Information Sciences, Middlesex University, London, NW4 4BT, United Kingdom.; School of Computing and Mathematics, Keele University, ST5 5BG, United Kingdom.

**Keywords:** fluctuation test, antibiotic resistance, fitness, rifampicin, nalidixic acid, mutagenesis

## Abstract

Evolution depends on mutations. For an individual genotype, the rate at which mutations arise is known to increase with various stressors (stress-induced mutagenesis – SIM) and decrease at high population density (density-associated mutation-rate plasticity – DAMP). We hypothesised that these two forms of mutation rate plasticity would have opposing effects across a nutrient gradient. Here we test this hypothesis, culturing *Escherichia coli* bacteria in increasingly rich media. We distinguish an increase in mutation rate with added nutrients through SIM (dependent on error-prone polymerases Pol IV and Pol V) and an opposing effect of DAMP (dependent on MutT, which removes oxidised G nucleotides). The combination of DAMP and SIM result in a mutation rate minimum at intermediate nutrient levels (which can support 7×10^8^ cells ml^−1^). These findings demonstrate a strikingly close and nuanced relationship of ecological factors – stress and population density – with mutation, the fuel of all evolution.

## Introduction

How and why the rate of spontaneous genetic mutation varies is a fundamental and enduring biological issue (Lynch et al 2016). Mutation rate can vary both among species (Sung et al 2012) and within a genotype (Maharjan and Ferenci 2017b). Intra-genotypic variation can depend upon stressful environmental conditions such as nutrient limitation, growth-rate reduction, high osmotic pressure, low pH, extreme shifts in temperature or various DNA damaging agents (Foster 2007, Galhardo et al 2007, MacLean et al 2013). In these environments cells induce stress responses that can increase mutation rates, typically via up-regulation of error-prone polymerases Pol IV and Pol V (Al Mamun et al 2012) – a phenomenon known as stress-induced mutagenesis (SIM).

Recently, we found that across microbes, the mutation rate of a particular genotype critically depends on the density to which the population grows (that is, the carrying capacity of the environment divided by its volume) (Krašovec et al 2017). In this so-called Density-Associated Mutation-rate Plasticity (DAMP), bacterial and yeast populations show a power law (log-log linear) reduction in mutation rate with *D* when grown in a defined minimal medium with glucose as the sole carbon source (Krašovec et al 2014, Krašovec et al 2017). DAMP and SIM modify mutation rates in *Escherichia coli* via different genetic pathways. DAMP requires a Nudix hydrolase protein, whose primary role is degrading highly mutagenic 8-oxo-dGTP (Michaels and Miller 1992), while error-prone polymerases such as Pol IV is not involved in DAMP (Krašovec et al 2017). Differences in the underlying mechanism and the fact that the most dense populations, experiencing the highest stress, show the lowest mutation rates, suggest that DAMP is not obviously associated with stress.

Growth in minimal medium on a single carbon source does not, however, reflect the environmental complexity or range of population densities experienced by many species. *E. coli* population density in host environments varies over 5 orders of magnitude among host species, and can be higher than 10^9^ colony forming units per gram of faeces (reviewed in (Tenaillon et al 2010)). As the highest population densities, with the greatest competition, rely on high nutrient availability, we reasoned that the addition of nutrients to minimal nutrient environments could indirectly increase both population densities and the level of stress. We therefore hypothesised that effects of density and stress on mutation rates, DAMP and SIM respectively, will act in opposition to one another across such a nutrient gradient – DAMP decreasing mutation rate and SIM increasing it as nutrients and population density increase.

Here we test this hypothesis by determining *E. coli* mutation rates across a nutrient gradient, while genetically manipulating DAMP and SIM independently. As hypothesised, we identify genetically separable and opposed associations of mutation rate with nutrient availability – a negative association requiring *mutT* (DAMP) and a positive association requiring polymerases IV and V (*dinB* and *umuC* respectively; SIM). We find that these associations combine to minimise average mutation rates in environments with intermediate nutrient availability and population density.

## Results

We assayed mutation rates to rifampicin resistance using fluctuation tests in *E. coli* K-12 MG1655 grown across a gradient of nutrient availability: a range of concentrations (1-90% *v/v)* of lysogeny broth (LB) mixed with Davis minimal (DM) medium (LB/DM). We find that the relationship of mutation rate to LB concentration is non-linear (Fig. 1, likelihood ratio test of a quadratic effect of log nutrient availability on log mutation rate: *N*=97, *LR*_8,7_=105, *P*=1.2×10^−24^, model S-I in Supplementary Information). Mutation rate to rifampicin resistance decreases as LB/DM is increased from 1% to 10% LB (increasing final population density, *D*, from 1×10^8^ to 7×10^8^ cells ml^−1^ Fig. S2-S3). This is comparable to DAMP in DM with glucose (Krašovec et al 2014, Krašovec et al 2017). However, mutation rate starts to increase in richer media, with 90% LB reaching similar or higher mutation rates than in 1% LB.

**Figure 1.**
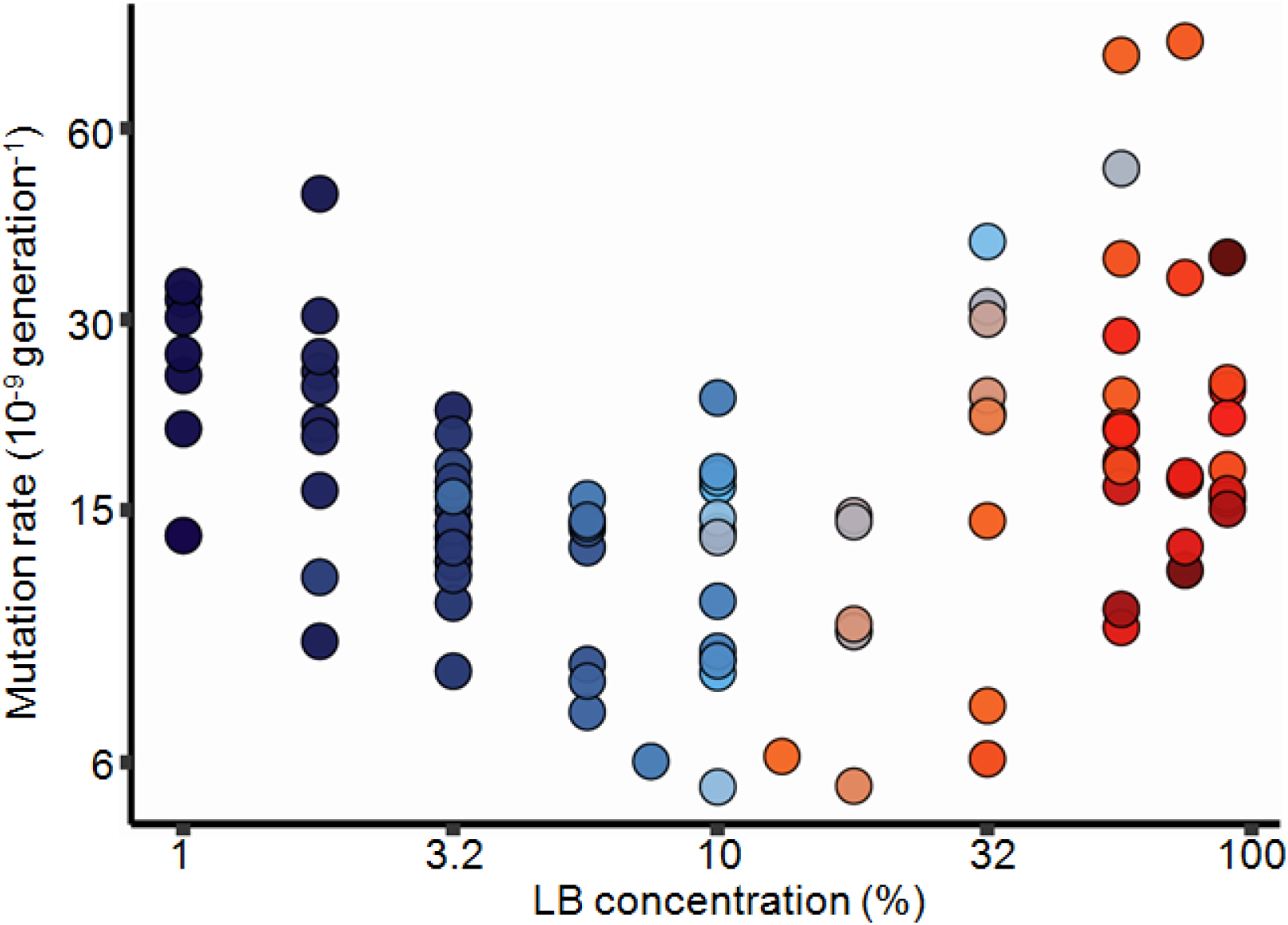
Effect of nutrient availability on mutation rate to rifampicin resistance in wild-type *E. coli* K-12 MG1655 (see model S-I in Supplementary Information *N*=97). Cells were grown in Davis minimal medium mixed with 1% to 90% of lysogeny broth (LB) medium. Colours represent a range of population densities from 4.5×10^7^ (dark blue) to 3.3×10^9^ (dark red, see Fig. S1 for details of the scale). See Figure S3 for a plot of mutation rate directly against population density and S9 for mutation rates co-estimated with the relative fitness of resistant mutants. Note the non-linear axes.

We next asked whether the increase in mutation rate at higher nutrient availability is genetically separable from the decrease in mutation rate due to DAMP. DAMP in *E. coli* requires the 8-oxo-dGTP diphosphatase MutT protein, meaning that, in minimal medium with glucose, the mutation rate in a Δ*mutT* mutant does not decrease with increased nutrient concentration (Krašovec et al 2017). We therefore performed fluctuation tests to nalidixic acid resistance in LB/DM with a Δ*mutT* mutant. We find that in LB/DM, as in DM with glucose, mutation rate in Δ*mutT* shows no relationship with increased nutrients or population density below 10% LB (Fig. 2, Model S-II in Supplementary Information). However, even more clearly than in the wild-types (both MG1655 [Fig. 1] and the immediate parent of the Δ*mutT* mutant [Fig. S6]), mutation rate of the Δ*mutT* mutant increases with the nutrient availability above 10% LB (population density of ~1 ×10^9^ cells ml^−1^). The only other *E. coli* mutant reported not to exhibit DAMP is *E. coli* K-12 MG1655 Δ*luxS* (Krašovec et al 2014). However, this mutant’s deficiency in DAMP is functionally complemented by added aspartate (Krašovec et al 2014), and LB is a medium rich in amino acids (Sezonov et al 2007). If variation in mutation rate at 10% LB/DM and below is the same phenomenon as DAMP, we expect this mutant strain to behave more similarly to wild-type than the Δ*mutT* mutant. We find that the Δ*luxS* mutant’s mutation rate is indistinguishable from the wild-type MG1655 across LB/DM environments (Fig. S6, *N*=167, likelihood ratio tests of the *luxS* deletion on the interaction between quadratic response of mutation rate to LB concentration and genotype [*N* = 167, *LR*_11,12_=2, *P*=0.16], and on the fixed effect of genotype *[N =* 167, *LR*_11,10_=1.4, *P=0.24*]).

**Figure 2.**
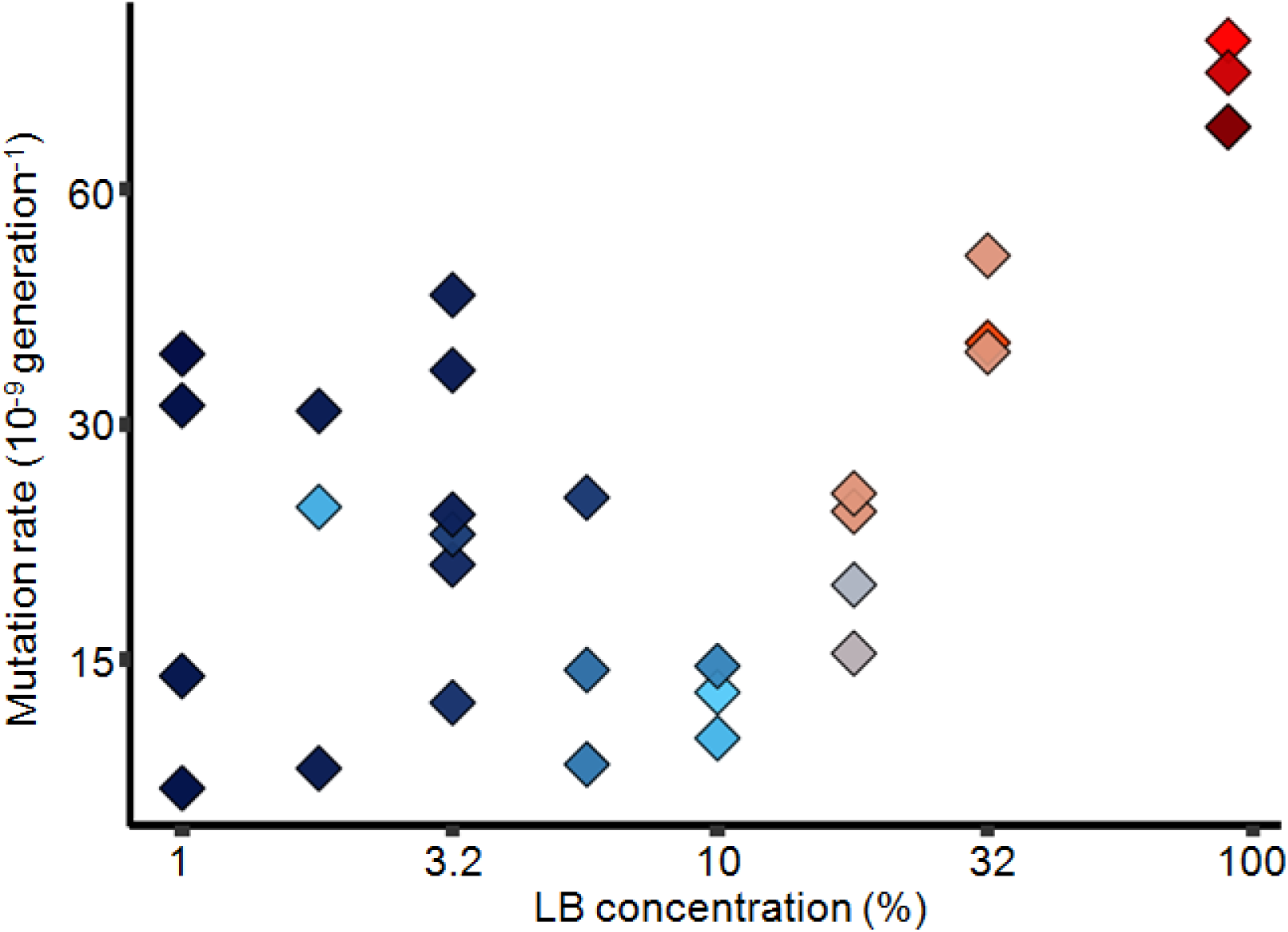
Effect of nutrient availability on mutation rate to nalidixic acid resistance in cells without DAMP (Δ*mutT, N*=30). Cells were grown in Davis minimal medium mixed with 1% to 90% of lysogeny broth (LB) medium. Colours represent a range of population densities from 4.5×10^7^ to 4.5×10^9^ (see Fig. S1 for details of the colour scale). See Fig. S5 for a plot of mutation rate directly against population density and S10 for mutation rates co-estimated with the relative fitness of resistant mutants. Note the non-linear axes.

The fact that mutation rate increases at high LB concentrations in a Δ*mutT* mutant (Fig. 2), where DAMP is absent, suggests that high nutrient availability increases mutation rate via a DAMP-independent mechanism. We hypothesised that higher nutrient concentrations increase the level of stress (e.g. by promoting competition), thereby causing error-prone polymerases Pol IV and Pol V (coded by *dinB* and *umuC* respectively) to increase the mutation rate at very high LB concentrations. We tested this hypothesis by estimating mutation rates to rifampicin resistance in *E. coli* Δ*dinB* and Δ*umuC* growing in LB/DM. We find that, unlike *E. coli* MG1655 (Fig. 1) and Δ*mutT* (Fig. 2), mutation rates of the Δ*dinB* and Δ*umuC* deletants (Fig. 3, Model S-III in Supplementary Information) decrease with increasing nutrients above 10% LB (above population densities of 7×10^8^ cells ml^−1^, Fig. S7). This continued decrease indicates that these polymerases are required for the rise in mutation rates as nutrients increase, and that DAMP continues to affect mutation rates at high nutrient levels.

**Figure 3.**
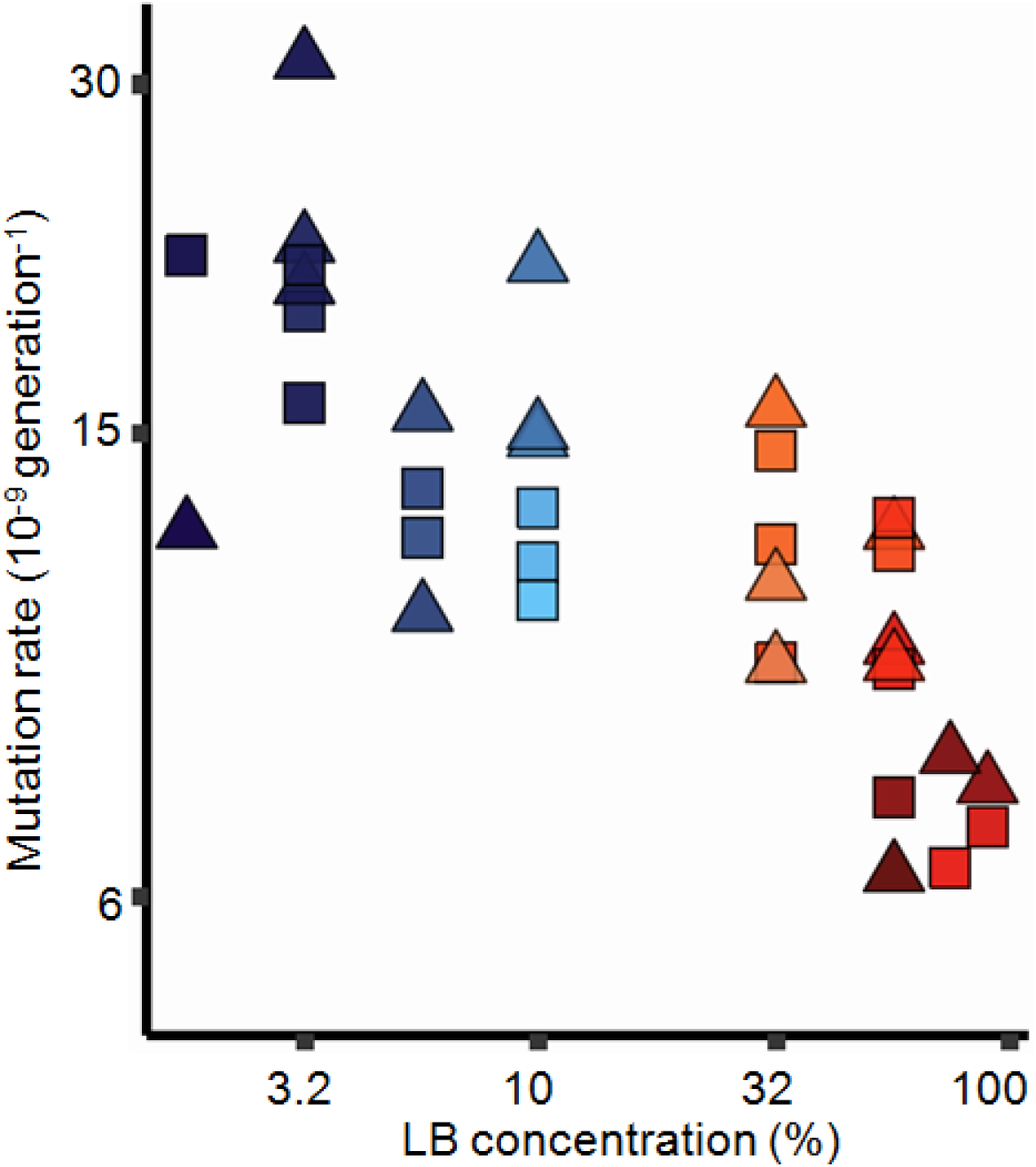
Effect of nutrient availability on mutation rates to rifampicin resistance in cells without error-prone polymerases Pol IV (Δ*dinB, N*=18, triangles) and Pol V (Δ*umuC, N*=18, squares). Cells were grown in Davis minimal medium mixed with 1% to 90% of lysogeny broth (LB) medium. The mutation rates of the two strains are not distinguishable (likelihood ratio test of the effect of genotype [*N*=36, *LR*_9_,_8_=2.6, *P*=0.11]). There is no evidence of a non-linear relationship of log mutation rate with nutrient availability (likelihood ratio test of a quadratic effect [*N*=36, *LR*_8_,7=0.15, *P*=0.70]), but there is a highly significant linear effect of nutrient availability (likelihood ratio test of linear effect [*N*=36, *LR*_7_,6=45, *P*=1.6×10^−11^], see model S-IV in Supplementary Information). Colours represent a range of Δ*dinB* and Δ*umuC* population densities, 4.8×10^7^ - 3.5×10^9^ and 5.6×10^7^ - 3.2×10^9^, respectively (see also Fig. S1). See Figure S7 for an equivalent plot using population density and S11 for mutation rates co-estimated with the relative fitness of resistant mutants. Note the non-linear axes.

The fitness effects of resistance mutations are known to be variable among nonselective environments, particularly for rifampicin (Maharjan and Ferenci 2017a). This variation has the potential to give artefactual differences in mutation rates among environments. We therefore estimated the fitness effects of resistance mutations in the fluctuation tests reported in Figs. 1-3 (Fig. S8). As expected, resistance mutations were, on average, somewhat deleterious across nutrient environments for both rifamipcin and nalidixic acid. Surprisingly, the average effect was least deleterious in intermediate nutrient environments. This suggests that, if anything, mutation rates in intermediate nutrient environments are over-estimated, relative to high and low nutrient environments. Thus, the results reported in Figs. 1-3 are robust to environmentally-dependent fitness effects of resistance mutations (Fig. S9-11).

## Discussion

Our previous work on density associated mutation rate plasticity (Krašovec et al 2017) contained a paradox. In the laboratory, *E. coli* displays a substantial and highly significant decrease in mutation rate with population density (DAMP, much more so than related bacteria – *Pseudomonas aeruginosa* PAO1); yet, in the published literature, of all 56 species with appropriate data (including *P. aeruginosa)*, the one with least negative association was *E. coli*. Here we have resolved that paradox. We have shown how two mechanistically independent plastic processes act on the mutation rate: DAMP – apparent at lower population densities (< 7×10^8^ cells ml^−1^), causing mutation rate to *decrease* with nutrient concentration; SIM – apparent at higher population densities (> 1×10^9^ cells ml^−1^), causing mutation rate to *increase* with nutrient concentration. *E. coli* is the organism whose mutation rate has, across the last 75 years of literature, been measured across the broadest range of population densities (7.50×10^6^ to 8.85×10^9^ cells ml^−1^) (Krašovec et al 2017). This means that, like Fig. 1 (and Fig. S3), the published literature uses a range of media and shows a minimum in *E. coli’s* mutation rate at around 7×10^8^ cells ml^−1^ (Fig. 4). This explains why attempting to fit a linear trend to this data does not yield a steep negative relationship (Krašovec et al 2017). It is also consistent with DAMP acting at low population densities and SIM at high densities across diverse published studies (144 individual estimates across 18 studies), as we find here in a single, controlled study.

**Figure 4.**
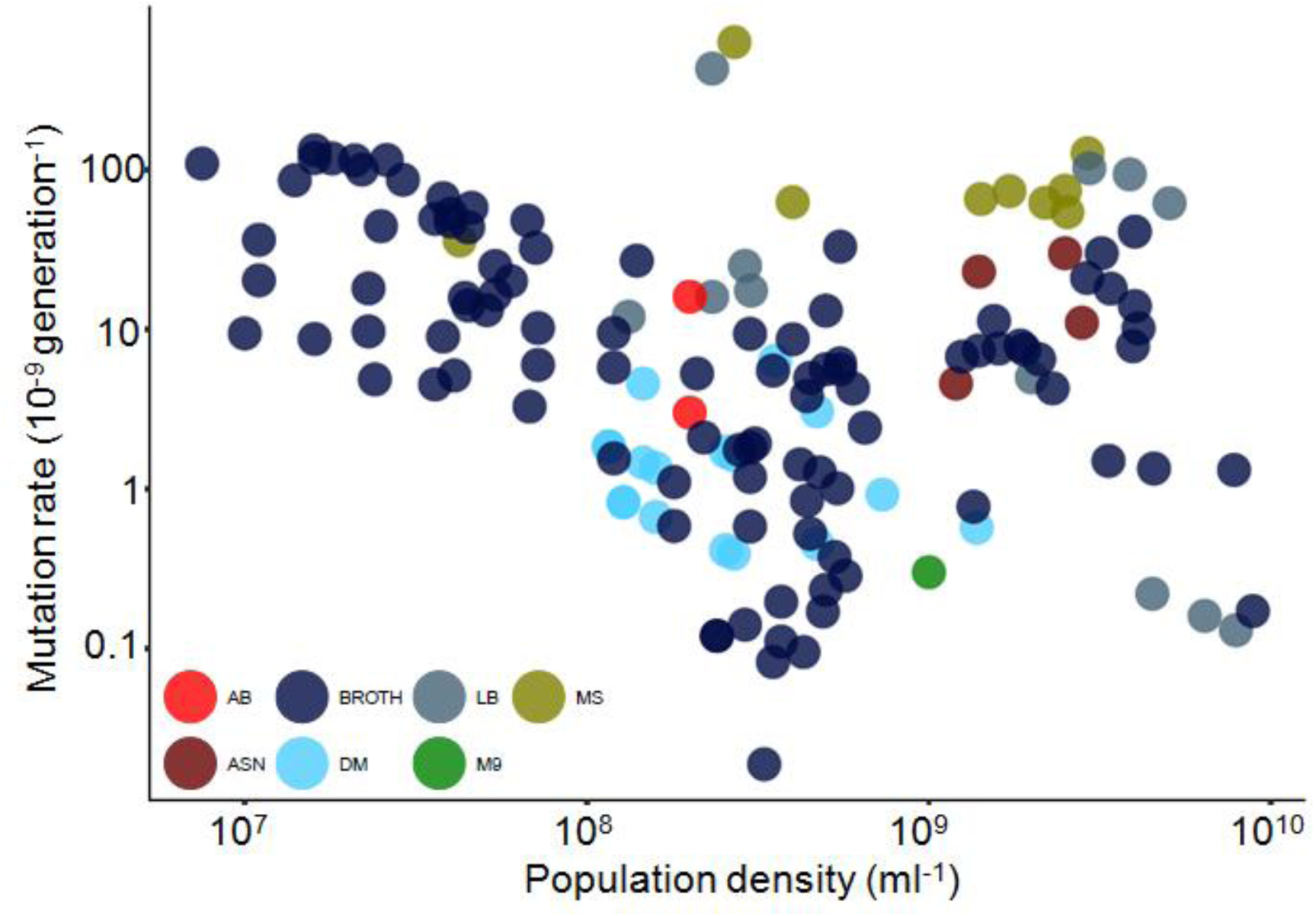
Distribution of *E. coli* mutation rates published from 1943 to 2017 (re-plotted from reference (Krašovec et al 2017), *N*=144). Abbreviations: AB - minimal medium with various carbon sources, ASN - asparagine glucose synthetic medium, BROTH-rich nutrient medium, DM - Davis minimal medium with various carbon sources, LB-rich lysogeny broth medium, M9-minimal medium with various sugars, and MS-minimal salt medium with various sugars. These data also include a range of strains and phenotypic markers (see Supplementary Data File). Note the non-linear axes.

It seems likely *a priori* that the dynamics of population growth and cell division, which differ among nutrient environments, are involved in the mutation rate changes observed here. For instance, environmental differences that affect growth rate will in turn affect ploidy (Pecoraro et al 2011), which can affect mutation rate estimates (Sun et al 2017). Therefore, we cannot exclude the possibility that the effects of either DAMP or SIM on mutation rate considered here are mediated by some aspect(s) of the culture cycle that differ across different nutrient environments. Such dynamics are largely inaccessible to fluctuation tests, as used here, or indeed other standard methods of assaying mutation rate (Foster 2006) that consider at least one full population growth cycle. It may become possible to assay such changes in the future with single-cell mutation monitoring approaches (Elez et al 2010, Uphoff et al 2016).

Variation in mutation rate among members of a population can itself provide evolutionary advantages (Alexander et al 2017), and modulating mutation rate in response to the environment could hypothetically allow organisms to optimise their rate of adaptation (Belavkin et al 2016). Such ‘optimal’ variation involves minimising mutation rates at high fitness, but allowing them to increase away from fitness peaks. However, environmental cues do not give direct information about an individual’s fitness. An organism may just receive information about the levels of particular molecules in the environment. These could give information, for instance, about availability of a particular nutrient or about population density, and that information may be linked to mechanisms involved in the plastic control of mutation rate, e.g. (Krašovec et al 2014). But a cue indicative of high population density could be an indicator of an individual having high fitness (if it is part of a successful clone), or high competition and therefore low individual fitness. Similarly, un-utilised nutrients may be indicative of a benign environment and therefore high fitness, or of a clone that has been unable to utilise resources and therefore low individual fitness. Organisms receive many environmental cues and may therefore be able to parse them to give a clear picture of the competitive environment, enabling appropriate responses (Cornforth and Foster 2013). How far this may occur in terms of mutation rate is unclear, and adaptive explanations for the existence of SIM are probably unnecessary, given more direct and/or non-adaptive explanations (Lynch et al 2016, MacLean et al 2013). Nonetheless, it is reasonable to speculate about the evolutionary effects of these plastic mutation rate traits, whatever their origins, either evolutionarily or mechanistically in terms of environmental cues. The effect of minimising mutation rate in intermediate nutrient (and population density) environments (Figs. 1-2, S3 and S5) may be to minimise evolutionary change for organisms that are doing well in a relatively benign nutrient environment, but without excessive competition, which is potentially advantageous (Belavkin et al 2016).

Without clearer evidence around the evolution of these traits and their effects on evolution beyond fluctuation tests, any reasoning about their adaptive effects remains speculative. Nonetheless, population density and nutrient availability are focal points of microbial ecological competition (Hoek et al 2016, Oliveira et al 2015). Microbes have evolved numerous strategies to sense and increase acquisition of resources (Hibbing et al 2010), and they possess efficient ways of sensing population density (Xavier and Bassler 2005). A threshold population density (known as quorum) is often required to regulate a diverse array of physiological activities (Walters and Sperandio 2006), many of which promote stress tolerance (Williams et al 2007). The fact that mutation rate too responds in a complex way to such environmental factors suggests that, for the *de novo* evolution of traits such as antibiotic resistance, ecological circumstances and evolutionary outcomes are tightly linked.

## Acknowledgments

We thank Karina B. Xavier for *E. coli* MG1655. We thank César Aguilar, Christina Moon, Colin Russell and Sebastian Wielgoss for providing data. We thank Mark Foster and John Parfitt for technical assistance.

## Materials and Methods

### Strains used in this study

*Escherichia coli* K-12 strain KX1228 (Δ*luxS)* was derived from the wild-type K-12 MG1655 *(luxS^+^)* (Xavier and Bassler 2005). *E. coli ΔmutT* mutant is part of Keio collection designated as JW0097-1 (F-, Δ*(araD-araB)567, ΔlacZ4787(∷*rrn*B-3)*, λ-, Δ*mutT790∷kan, rph-1, Δ(rhaD-rhaB)568, hsdR514). E. coli* Δ*dinB* mutant is part of Keio collection designated as JW0221-1 (F-, Δ*(araD-araB)567, ΔlacZ4787(∷*rrn*B-3)*, λ-, Δ*dinB749:*:kan, *rph-1*, Δ*(rhaD-rhaB)568, hsdR514. E. coli ΔumuC* mutant is part of Keio collection designated as JW1173-1 (F-, Δ*(araD-araB)567, ΔlacZ4787*(∷rrnB-3), λ-, Δ*umuC773∷kan, rph-1*, Δ*(rhaD-rhaB)568, hsdR514)*. Parent of Keio collection is an *E. coli* strain BW25113 (F-, Δ*(araD-araB)567, ΔlacZ4787(∷*rrn*B-3)*, λ-, *rph-1*, Δ*(rhaD-rhaB)568, hsdR514*).

### Media

We used MilliQ water for all media. Strains were grown with shaking (250 rpm) at 37°C in lysogeny broth (LB) medium (10g NaCl, 5g yeast extract and 10g tryptone per litre [l^−1^]) mixed with Davis minimal (DM) medium (0.5g C_6_H5Na3O_7_×2H20, 1g (NH_4_)_2_SO4, 2g H_2_KO_4_P and 7g HK_2_O_4_P×3H_2_O l^−1^). 100mg L^−1^ MgSO_4_×7H_2_0 (406 μmol) and 4μg L^−1^ thiamine hydrochloride were added to DM after autoclaving. We used 1% to 90%LB with a content of Mg^2+^ ranging from 40 to 35.5 μmol respectively, assuming that LB contains on average 35 μmol l^−1^ of Mg^2+^ (Papp-Wallace and Maguire 2008). Selective tetrazolium arabinose agar (TA) medium (10g tryptone, 1g yeast extract, 5g NaCl, 3g arabinose and 0.05g 2,3,5-Triphenyl-tetrazolium chloride l^−1^) was supplemented with freshly prepared rifampicin (50μg ml^−1^) or nalidixic acid (30μg ml^−1^). For all cell dilutions sterile saline (8.5 g l^−1^ NaCl) was used. All media were solidified as necessary with 15 g l^−1^ of agar (Difco).

### Fluctuation tests

We did fluctuation tests with *Escherichia coli* as already explained (Krašovec et al 2014, Krašovec et al 2017). In short, strains were first inoculated from frozen stock and grown in liquid LB medium at 37°C and then transferred to nonselective liquid media (LB or DM with glucose) and allowed to grow overnight shaking at 37°C. *E. coli* cells were again diluted into fresh LB/DM, giving the initial population size (*N*_0_) of 2,373 (range 1.5×10^2^ – 1.3×10^4^). Various volumes (0.35–1 ml) of parallel cultures were grown to saturation for *24 hours* at 37°C in 96 deep-well plates. The position of each culture on a 96-deep-well polypropylene plate was chosen randomly. Final population size *(N_t_)* was determined by colony forming units (CFU) where appropriate dilution was plated on solid non-selective TA medium. Population density (*D*) was estimated was determined by two independent techniques using CFU and an ATP based assay: luminescence (LUM) was measured using a Promega GloMax luminometer and the Promega Bac-Titer Glo kit, according to manufacturer’s instructions. We measured luminescence of each culture 0.5 and 510 seconds after adding the Bac-Titer Glo reagent and calculated net luminescence as LUM = luminescence_510_s – luminescence_0.5_s. Each estimate of *D* and *N_t_* was averaged across 3 independent cultures. Evaporation (routinely monitored by weighing plate before and after 24h incubation) was accounted for in the *N_t_* value determined by CFU and was also used in statistical modelling as a variance covariate. We obtained the observed number of mutants resistant to rifampicin or nalidixic acid, *r*, by plating the entirety of remaining cultures onto solid selective TA medium (4.5cm plates in Figs. 1-2 and 9cm plates in Fig. 3) that allows spontaneous mutants to form colonies. Plates were incubated at 37°C and mutants were counted at the earliest possible time after plating. For rifampicin plates this was 44–48 hours, when nalidixic acid was used the incubation time was 68–72 hours.

For figures 1, 2 and 3 we used 13, 3 and 4 independent experimental blocks, respectively, carried out on different days. Within an experimental block multiple 96-well plates were used. Any individual mutation rate estimate requires multiple parallel cultures, which were all carried out on a particular plate. For figures 1, 2 and 3 the median (with interquartile range) of parallel cultures used per mutation rate estimate was 16 (21–16), 16 (16–16) and 16 (16–16), respectively.

### Estimation of mutation rates

To estimate number of mutational events, *m*, from the observed number of mutants we employed the Ma-Sandri-Sarkar maximum-likelihood method implemented by the FALCOR web tool (Hall et al 2009). The mutation rate per cell per generation is calculated as *m* divided by the final population size, *N_t_*, determined by CFU. This approach does not account for potentially important issues that may affect mutation rate estimates. Crucially, if there is a cost to carrying a resistance allele in the fluctuation test environment, this can result in an under-estimation of the mutation rate. This issue can be corrected for by co-estimating the average fitness effect of resistance mutations with the mutation number of mutational events (Zheng 2005). In addition, variation in *N_t_* may also affect estimates and may also be accounted for (Ycart and Veziris 2014). We therefore co-estimated mutation rates and fitness effects, accounting for variability in *N_t_* using the flan package in R (Adrien et al 2017), also setting the Winsorization parameter to remove the effects of ‘jackpots’ with un-countably large numbers of mutants (greater than 150 on 4.5cm plates and greater than 1000 on 9cm plates). Since the estimated fitnesses (Fig. S8) tend to reinforce the patterns seen in Fig. 1-3, we report the results of the simpler and more widely used calculations in the main text as being more conservative.

### Statistical analysis

All statistical analysis was executed in R v3.3.1 (R Core Team 2016) and nlme v3.1 packages for linear mixed effects modelling (Pinheiro and Bates 2000). This enabled the inclusion within the same model of experimental factors (fixed effects), blocking effects (random effects) and factors affecting variance (giving heteroscedasticity) as described in the Supplementary Information. In all cases log_2_ mutation rates were used. Details of models and their fitting are given in the Supplementary Information: diagnostic plots in Supplementary figures S12-S14, ANOVA tables for each model are given in Supplementary Tables S3-S5. Code and data to reproduce the main text figures are given in the accompanying R script, and Supplementary Data File respectively. The content of the Supplementary Data File is explained in Supplementary Table S2.

**Author contributions:**
RK and CGK designed experiments carried out by RK and analysed by RK and CGK. All authors contributed to scientific direction of the project. The paper was written by RK and CGK incorporating contributions from all other authors.

